# A WIDESPREAD PICORNAVIRUS AFFECTS THE HAEMOCYTES OF THE NOBLE PEN SHELL (*PINNA NOBILIS*) LEADING TO IMMUNOSUPPRESSION

**DOI:** 10.1101/2023.11.05.565683

**Authors:** Francesca Carella, Patricia Prado, Gionata De Vico, Dušan Palić, Grazia Villari, José Rafael García-March, José Tena-Medialdea, Emilio Cortés Melendreras, Francisca Giménez- Casalduero, Marco Sigovini, Serena Aceto

## Abstract

The widespread mass mortality of the noble pen shell (*Pinna nobilis*) has occurred in several Mediterranean countries in the past seven years. Single-stranded RNA virus affecting immune cells and leading to immune disfunction have been widely reported in human and animal species. Here we present data linking *P. nobilis* mass mortality events (MMEs) to haemocyte picornavirus (PV) infection. This study was performed on 30 specimens, from wild and captive populations. We sampled *P. nobilis* from two regions of Spain, Catalonia [24 animals] and Murcia [two animals]), and one region in Italy (Venice [four animals]). The low number of analyzed specimens was due to the scarcity of remaining individuals in the Mediterranean Sea. In 100% of our samples, ultrastructure revealed the presence of a virus (20nm diameter), capable of replicating within granulocytes and hyalinocytes, leading to the accumulation of complex vesicles of different dimensions within the cytoplasm. As the PV infection progressed, dead haemocytes, infectious exosomes, and budding of extracellular vesicles were visible, along with endocytic vesicles entering other cells. The THC (total haemocyte count) values observed in both captive (eight animals) (3.5 x 10^4^ - 1.60 x10^5^ ml^-1^ cells) and wild animals (14 samples) (1.90 - 2.42 x10^5^ ml^-1^ cells) were lower than those reported before MMEs. Sequencing of *P. nobilis* (six animals) haemocyte cDNA libraries revealed the presence of two main sequences of *Picornavirales*, family *Marnaviridae*. The highest number of reads belonged to animals that exhibited active replication phases and abundant viral particles from Trasmission Electron Microscopy (TEM) observations. These sequences correspond to the genus *Sogarnavirus* - a picornavirus identified in the marine diatom *Chaetoceros tenuissimus* (named *C. tenuissimus* RNA virus type II). Real time PCR performed on the two most abundant RNA viruses previously identified by *in silico* analysis revealed positive results only for the sequences similar to *C. tenuissimus* RNA virus. These results may not be considered conclusive of picornavirus identification in noble pen shell haemocytes, and require further studies. Our findings suggest that picornavirus infection likely causes immunosuppression, making individuals prone to opportunistic infections which is a potential cause for the MMEs observed in the Mediterranean.

## 1. Introduction

Since June 2016, thousands of individuals of the noble pen shell *Pinna nobilis* have died in the Mediterranean due to an extensive mass mortality event (MME) (Carella et al., 2019; Kersting et al., 2019; Zotou et al., 2020; Carella et al., 2020; Lattos et al., 2021). The wide geographic range of the phenomenon makes the mass mortality of the species one of the largest known marine wildlife epizootics mortality events to date in the Mediterranean Sea (Kersting et al., 2019; Katsanevakis et al., 2022). The event was first recorded along hundreds of kilometers of the southeastern coast of the Iberian Peninsula (Darriba et al., 2016). Mass mortalities were subsequently observed in the north-western Mediterranean (French and Italian coasts) and soon after in Greece, Cyprus, Turkey, Algeria, Tunisia, Morocco, Albania, and Croatia (Carella et al., 2019; Kersting et al., 2019; Zotou et al., 2020; Carella et al., 2020; Lattos et al., 2021). The cause of this MME has been attributed to different pathogens, particularly *Haplosporidium pinnae* and *Mycobacterium* spp. (Catanese et al., 2018; Carella et al., 2023a). The presence of a great variety of pathogens associated with MMEs, including several potentially opportunistic, suggests that disease pathogenesis leading to animal mortality may have other unidentified causes (Carella et al., 2020, 2023a).

The order *Picornavirales* comprises positive-strand RNA viruses ranging between 7,000 and 12,500 nt in length (Le Gall et al., 2008). Within this order, the family *Picornaviridae* comprises 29 genera (https://ictv.global/report/chapter/picornaviridae/picornaviridae) of picornaviruses (PV) (Wang-Shick 2017). Picornaviruses are small, icosahedral viruses with single-stranded, highly diverse positive-sense RNA genomes, associated with mild to severe diseases in vertebrates, invertebrates, and plants (Whitton et al., 2005; Stanway, 1990; Gustavsen et al., 2014). The family *Picornaviridae* includes some of the most important groups in the development of virology, comprising poliovirus, rhinovirus, and hepatitis A virus (Cifuente et al., 2019; Whitton et al., 2005). Virions consist of a capsid, with no envelope, surrounding a core of ssRNA, measuring from 22-33 nm in diameter (Zell et al., 2017; Baltimore, 1971; Le Gall et al., 2008). Next Generation Sequencing (NGS) has significantly expanded the order *Picornavirales* in recent years, through the identification of previously unknown viruses found associated with various taxa, including aquatic vertebrates and invertebrates (Arcier et al., 1999; Tang et al., 2005; Kapoor et al., 2008; Shi et al., 2016; Kim et al., 2017; Liu et al., 2021). The replication cycle takes place in the cytosol in tight association with reorganized cytoplasmic membranous structures (Below et al., 2012; Harak and Lohmann, 2015). Infection with PV can induce numerous changes in infected cells, with the massive accumulation of cytosolic double-membraned vesicles (DMVs) and replication organelles, possibly being the most important changes (Dales et al., 1965; Hsu et al., 2010; Suhy et al., 2000). Picornaviruses target intracellular membranes to generate complex membrane rearrangements of host organelles, such as the endoplasmic reticulum (ER), mitochondria, or endolysosomes (Cottam et al., 2009; Yang et al., 2020). Host intracellular membranes contain molecules, lipids and proteins, that serve as vehicles for intercellular communication in various (patho)physiological processes (Welsh et al., 2007). The membrane may provide optimal platforms for viral RNA synthesis by concentrating viral replicative proteins and relevant host factors, as well as hiding replication intermediates, contributing to the evasion of host innate immune sensors (Den Boom et al., 2010).

Here we report the discovery of a previously undescribed picorna-*like* virus infecting the immune cells of the noble pen shell *Pinna nobilis* sampled between 2021-2023 in different regions of Spain and Italy - where mortality events have been reported. Thirty specimens were analyzed using Transmission Electron Microscopy (TEM) to describe the morphology and self-assembly of virions within the haemocyte cytoplasm and major structural transformations occurring in infected host cells. NGS Illumina sequencing coupled to qPCR was performed to describe the identified virus and part of its genome.

## 2. Material and methods

### 2.1 Haemolymph collection and Investigation of viral agents in haemocytes

Due to its status as an Endangered species, sampling of *P. nobilis* was carried out under the permission of regional and national authorities for animal welfare (for Spain, Generalitat de Catalunya Identificació de l’expedient: SF/0003/23; Bank of species of the Mar Menor INF/2020/0017 promoted by the General Directorate of the Mar Menor and the University of Murcia; for Italy, Prot. MATTM 0016478 05/03/2020).A total of 30 animals were included in the study, collected from July 2021 to May 2023. Sampling was performed on natural populations along the Mediterranean coasts of Italy (n = four) and Spain (n = 18) and from animals maintained in captivity in two facilities in Spain (n = eight). The natural sites in Spain and Italy were selected according to the presence of remaining populations of *P. nobilis* although with the occurrence of mortality outbreaks (**Figure 1)**.

**Figure 1.**
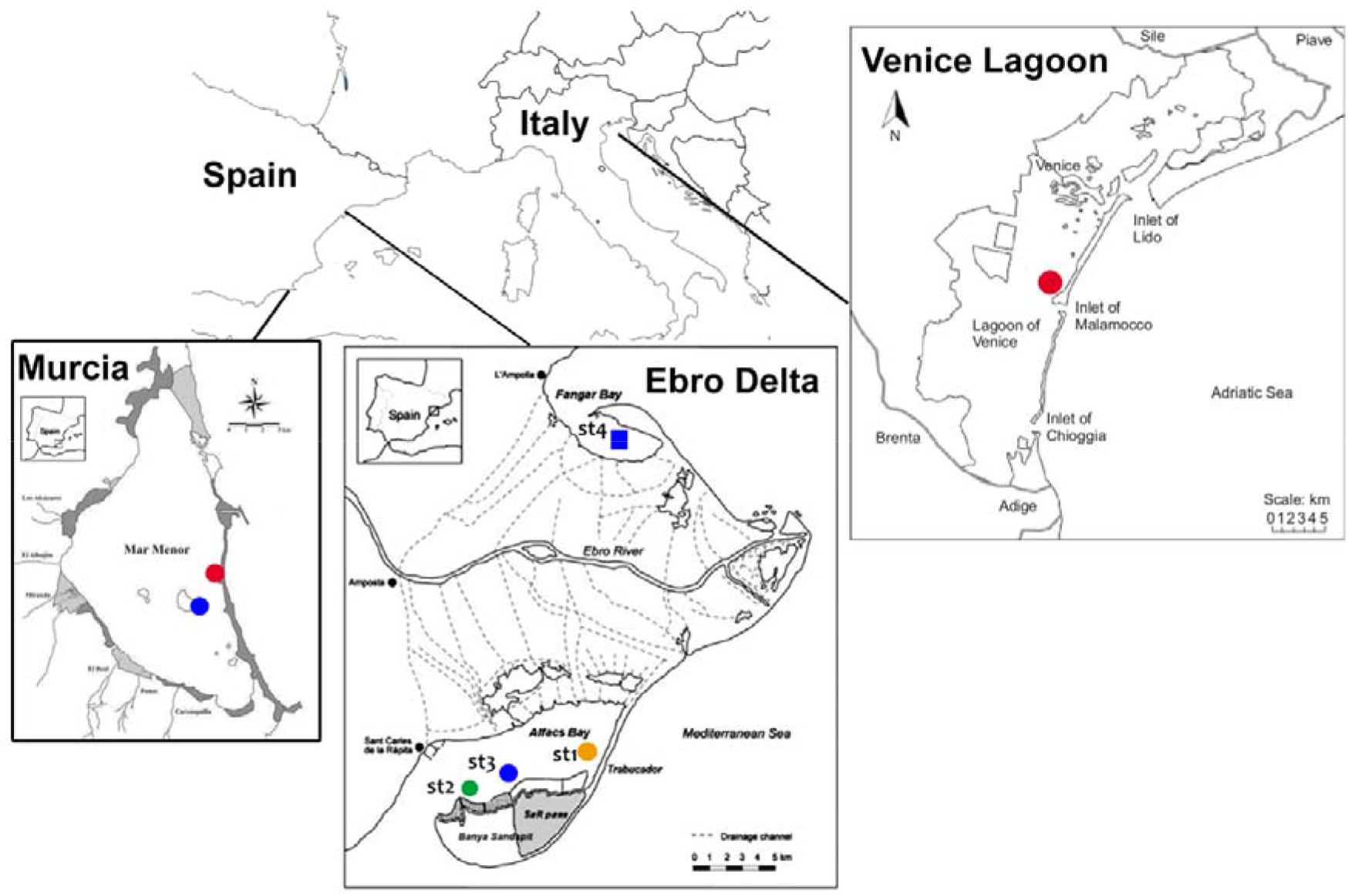
Map of the study areas where noble pen shell *Pinna nobilis* haemocytes were sampled in 2021-2023 in Murcia Baron (in blue) and La Manga (in red), Ebro Delta (site 1-4) and Venice lagoon (Ottagono Alberoni).

In Catalonia (Spain), samples were collected from two estuarine populations in the Ebro Delta, where residual populations still exist: the Fangar Bay (northern hemidelta) and the Alfacs Bay (southern hemidelta) distributed in 4 different monitored areas (24). Mortality was firstly observed in 2018 leaving only few individuals by 2021 (Prado et al. 2021). The mortality rate has progressively increasing from 33.5% in September 2021 to 75% in June 2023 (Prado pers. comm). Some of the remaining animals from this area are maintained in the aquarium facilities of the Instituto de Investigación en Medio Ambiente y Ciencia Marina (IMEDMAR) of the Universidad Católica de Valencia (UCV), in Valencia, Spain, and were included in the study. Another remaining population in Spain is the one located at the Mar Menor lagoon (Murcia), in the southwestern part of the Mediterranean. Is this area, *P. nobilis* mortality was first observed in the summer 2016. Initially it was correlated to an environmental collapse that became critical in 2016, 2019 and 2021 and reduced the population in >99 % (Gimenez-Casalduero et al., 2020); for the present work, two surviving localities were sampled in the Baron and La Manga areas (Cortes-Melendreras et al. 2022); some animals (n = two) from this area were transferred to the seawater tanks of the University of Murcia Aquarium. Animals (n = six) had been kept in captivity at IMEDMAR and Murcia Aquarium facilities for 6 months to one year. Until recently, *P. nobilis* was widely distributed in the Venice lagoon (Italy), with the largest and densest colonies over marine tidal deltas and the outer part of central basins (Tagliapietra et al., 2009). Mortality onset was observed at the end of 2020, when the population decreased between 60-80% (unplublished data). The animals (n = four) were sampled near the Ottagono Aberoni island, about 1 km away from the Malamocco inlet (Sigovini pers. Comm.).

For each animal included in the study (from the field and in captivity), 2 ml of haemolymph were collected via a 23-gauge needle from the posterior adductor muscle as a non-destructive sampling. Presence of disease clinical signs were considered during sampling. Literature reports that outward signs of disease start with behavioral changes, including shell gaping, slow valve closure speed when touched, and mantle retraction into the valves from the edge of the shell (Carella et al., 2023a). These signs were difficult to recognize in animals in the field and maintained in captivity, since they mostly appear at very late stages of disease (Carella et al., 2023a). In Catalonia, in July 21 and July/August 2022 most of the animals of the study (15) displayed apparent slow valve closure. In Venice lagoon, in May 2023, animals displayed quick valve closure and visible mantle at the border of the valves instead. Nevertheless, two months later, remarkable mortality was observed in the sampling area. Animals maintained in the Murcia Aquarium and IMEDMAR-UCV did not shown apparent signs of disease, but in both the cases, the animals suffered sudden mortaly of up to 40-60% in the following 6 months, primarily attributed to the manipulation from artificial spawning attempts.

After the procedure, the animals were immediately relocated into the sea bed/tanks and had their health monitored over the following days. A volume of about 2 ml of haemolymph was collected for each animal. One ml of haemolymph aliquot was placed in 1 ml of RNA Later (Sigma-Aldrich, Italy) labelled tube placed on ice for molecular analyses, and the other half was preserved for ultrastructural analyses. A small haemolymph aliquot (20 µl) was placed on a haemocytometer (Burker chamber) to determine the total cell number (x10^5^/ml) and define the total haemocyte count (THC) (**Table 1**).

**Table 1.** List of collected samples of noble pen shell *P. nobilis* haemolymph in the Mediterranean Sea; n.a.*data not available. Number of the samples analyzed per years are indicated within parentheses; THC: Total Haemocyte Count

| Site | Month and year | Number of collected animals evaluated by TEM | Sample Type | Mean THC (ml-1) of collected animals over the years |
| --- | --- | --- | --- | --- |
| *Alfacs Site 1 Trabucador* | July 21<br>July 2022 | 6 | Haemolymph | (4 in 2021): $2.35 \pm 0.82$ cells $\times 10^5$<br>(2 in 2022): $1.90 \pm 0.33$ cells $\times 10^5$ |
| *Alfacs Site 2 Xiringuito* | July 21<br>July 2022 | 5 | Haemolymph | (3 in 2021): $2.42 \pm 0.42$ cells $\times 10^5$<br>(2 in 2022): $2.15 \pm 0.34$ cells $\times 10^5$ |
| *Alfacs Site 3 Barca* | July 2021<br>August 2022 | 4 | Haemolymph | (3 in 2021) $2.30 \pm 1.0$ cells $\times 10^5$<br>2022 n.a.* |
| *Fangar Site 4* | May 2023 | 3 | Haemolymph | n.a.* |
| *Murcia aquarium* | September 22 | 2 | Haemolymph | $1.60 \pm 0.22$ cells $\times 10^5$ |
| *Tanks of IMEDMAR-UCV* | July 21<br>May 2023 | 6 | Haemolymph | (4 in 2021) $7.9 \pm 0.3$ cells $\times 10^4$<br>(2 in 2023) $3.5 \pm 0.3$ cells $\times 10^4$ |
| Venice lagoon | May 2023 | 4 | Haemolymph | n.a.* |

### 2.2 Transmission Electron Microscopy (TEM)

For all the animals included in the study, part of the haemolymph was also fixed in glutaraldehyde 2.5% in PBS for 3 h. Cells were postfixed in 1% osmium tetroxide for 60 min and 0.25% uranyl acetate overnight. They were dehydrated through a graded ethanol series and embedded in EPON resin overnight at 37 °C, 1 day at 45 °C and 1 day at 60 °C. Ultrathin sections (80 nm) were cut parallel to the substrate and placed onto a 200-mesh copper grid. Bright-field TEM images were obtained on the dried sample by using a FEI TECNAI G2 200 kV s-twin microscope operating at 120 kV (Thermo Fisher Scientific, Waltham, USA). Digital images were acquired with an Olympus VELETA camera.

### 2.3. RNA Extraction, Sequencing, and Bioinformatic Analysis

Total RNA was extracted from the haemocytes of six samples of*. nobilis* collected from 3 sites of wild population in Alfacs Bay, Catalonia (**Table 2**) using the PureLink RNA Mini Kit (Thermo Fisher Scientific, Waltham, Massachusetts, USA). Following DNase treatment, RNA concentration and quality was measured using the Nanodrop 2000 spectrophotometer (Thermo Fisher Scientific). Strand-specific cDNA libraries were prepared, and Illumina sequencing (NovaSeq 6000, 150 bp paired-end) was carried out by Eurofins Genomics (Ebersberg, Germany). Raw reads underwent trimming and adapter clipping using Trimmomatic 0.4 (Bolger et al., 2014).

**Table 2.** Samples of noble pen shell *P. nobilis* used to extract RNA from haemocytes and raw data sequencing statistics.

| Sample ID | Collection place | Collection date | Sequenced reads | Sequenced bases |
| --- | --- | --- | --- | --- |
| Bar1 | Alfacs Bay site 3 | August 2022 | 25,189,100 | 7,556,730,000 |
| Bar2 | Alfacs Bay site 3 | August 2022 | 28,331,700 | 8,499,510,000 |
| Trab8a | Alfacs Bay site 1 | July 2022 | 27,304,600 | 8,191,380,000 |
| Trab8b | Alfacs Bay site 1 | July 2022 | 29,977,200 | 8,993,160,000 |
| Xir2a | Alfacs Bay site 2 | July 2022 | 27,649,600 | 8,294,880,000 |
| Xir5a | Alfacs Bay site 2 | July 2022 | 23,687,100 | 7,106,130,000 |

To identify RNA viruses eventually infecting the *P. nobilis* haemocytes, the trimmed, good-quality reads were aligned to the *P. nobilis* genome (ASM1616189v1) using Bowtie2 (Langmead & Salzberg, 2012), and the unmapped reads were extracted using Samtools (Danecek et al., 2021). A total of 33,097,808 paired unmapped reads were assembled using Trinity 2.12.0 (Grabherr et al., 2011), and the abundance estimation of contigs was performed with RSEM (Li & Dewey, 2012). The Trinity-assembled contigs (140,922) were used as queries in a BLASTX search against the viral protein database (https://ftp.ncbi.nlm.nih.gov/refseq/release/viral/, downloaded on 6 June 2023). To focus the annotation on RNA viruses, the assembled contigs were analyzed using Virsorter2 (Guo et al., 2021) by selecting only the classification of RNA viruses. Additionally, VirBot, an RNA viral contig detector for metagenomic data (Chen et al., 2023), was utilized with the sensitive option. The contigs identified as positive by both Virsorter2 and VirBot were subjected to analysis using ORFfinder (https://www.ncbi.nlm.nih.gov/orffinder/). The resulting ORFs were then subjected to a BLASTP search against the UniProtKB/Swissprot, RefSeq protein, and non-redundant protein sequences databases on 4 July 2023.

The predicted sequences of the RNA-dependent RNA polymerase (RdRp) protein encoded by the identified RNA viruses were used for BLASTP searches against the viral protein database, and the highest-scoring homologous sequences were downloaded. Multiple amino acid alignments of the selected RdRp sequences was performed using the Constraint-based Multiple Alignment Tool (COBALT, https://www.ncbi.nlm.nih.gov/tools/cobalt/re_cobalt.cgi). The best amino acid substitution model was searched, and the maximum likelihood tree was generated using the LG + G + I model and 500 bootstrap replicates using MEGAX (Kumar et al. 2018), resulting in a final dataset of 317 characters.

### 2.4 Quantitative real-time PCR to detect the virus presence in the *Pinna* haemocytes

Total RNA was extracted from the haemocytes of five *P. nobilis* samples collected from different areas in of Alfacs Bay, Catalonia (**Table 3**) as described before. RNA (500 ng) was reverse-transcribed using the Maxima First Strand cDNA Synthesis (Thermo Scientific). Specific primer pairs were designed to amplify the two most abundant RNA viruses identified by the *in silico* analysis (**Table 4**), and quantitative real-time PCR experiments were conducted. Amplification reactions were performed in a total volume of 10 μl using 5 μl of PowerUp SYBR Green Master Mix (Applied Biosystems), 0.2 μM of each specific primer and adjusted to 10 μl with distilled water. The reactions were conducted in technical duplicates, using the *P. nobilis Elongation Factor*-1 (*EF-1*) as reporter gene (Lattos et al., 2023; Huan et al., 2016) (**Table 4**). The amplification conditions were the following: 1 cycle for 2 min at 50°C; 1 cycle for 2 min at 95°C, 40 cycles of amplification at 95°C for 15 s, 60°C for 18 s and 72°C for 1 min. The reactions were conducted in the QuantStudio 3 Real-Time PCR System (Thermo Scientific). The cDNA of the samples identified as Trab8a and XIR2a (the same used in the RNA-seq experiment) were subjected to real-time PCR amplification to confirm the results of the *in silico* analysis, and two negative controls were run without cDNA.

**Table 3.** Samples of noble pen shell *P. nobilis* used to extract RNA from haemocytes used for the qPCR.

| Sample ID | Collection place | Collection date |
| --- | --- | --- |
| Bar6 | Alfacs Bay site 3 | August 2022 |
| Bar7 | Alfacs Bay site 3 | August 2022 |
| Trab3 | Alfacs Bay site 1 | July 2022 |
| Trab5 | Alfacs Bay site 1 | July 2022 |
| Xir6 | Alfacs Bay site 2 | July 2022 |

**Table 4.**
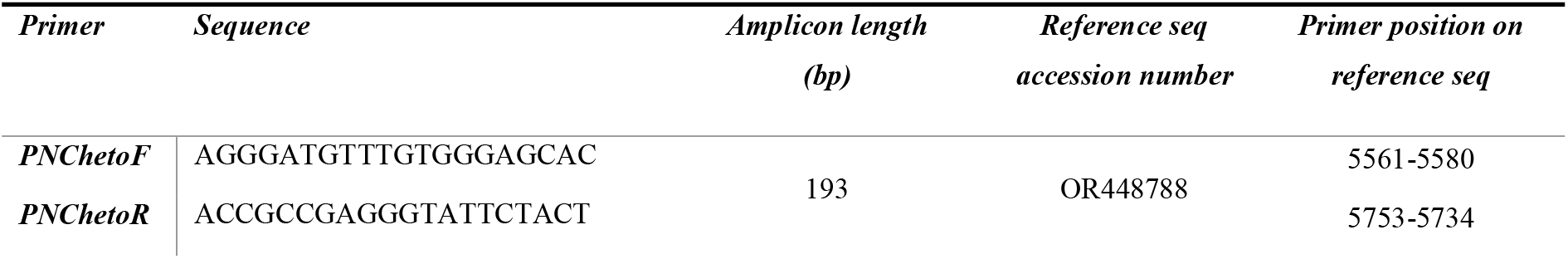
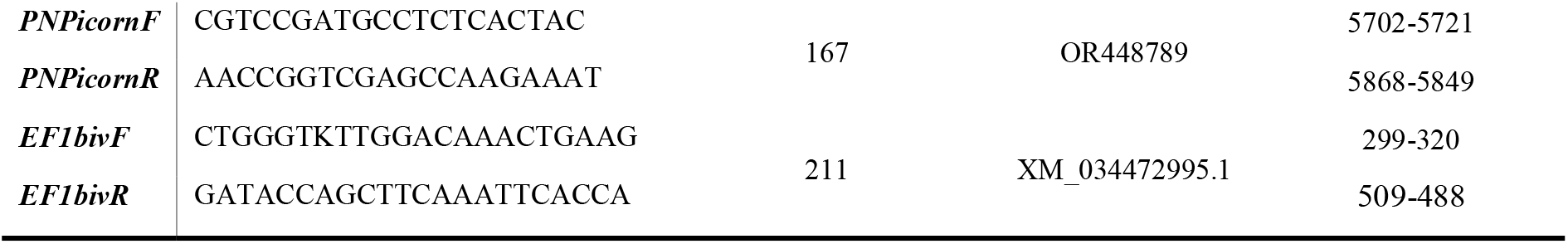
Primers used for the assessment of virus presence in noble pen shell *Pinna nobilis* haemocytes.

## 3. Results

### 3.1 Total Hemocyte Count, Ultrastructure of infected haemocytes and virus details

The THC values varied among individuals and areas over the years. Mean haemocyte count showed the lowest values from animals maintained in captivity in IMEDMAR-UCV and Murcia Aquarium (between 3.5 x 10^4^ - 1.60 x10^5^ ml^-1^ cells) compared with the natural population in Catalonia (1.90-2.42 x 10^5^ ml^-1.^ cells) (**Table 1**). The most represented cells were granulocytes (∼10 µm) displaying a small peripheral nucleus, with a small quantity of hyalinocytes (∼7 µm), presenting a large central nucleus and small cytoplasm. All the haemocytes collected from animals in all areas of Italy and Spain from 2021 to 2023 (100%) displayed the ultrastructural feature of viral infection with associated cell death. They all exhibited complex and unique membrane relocations displaying numerous cytoplasmic vesicles associated with damaged mitochondria, total lack of granules, presence of protein aggregates, and, in some cases, cellular buddings. Formation of invaginations occurred at the membrane of various organelles, including ER, endolysosomes, and mitochondria (**Figures 2-3**). Features of the viral infection at haemocyte level is represented in **Figure 2**.

**Figure 2.**
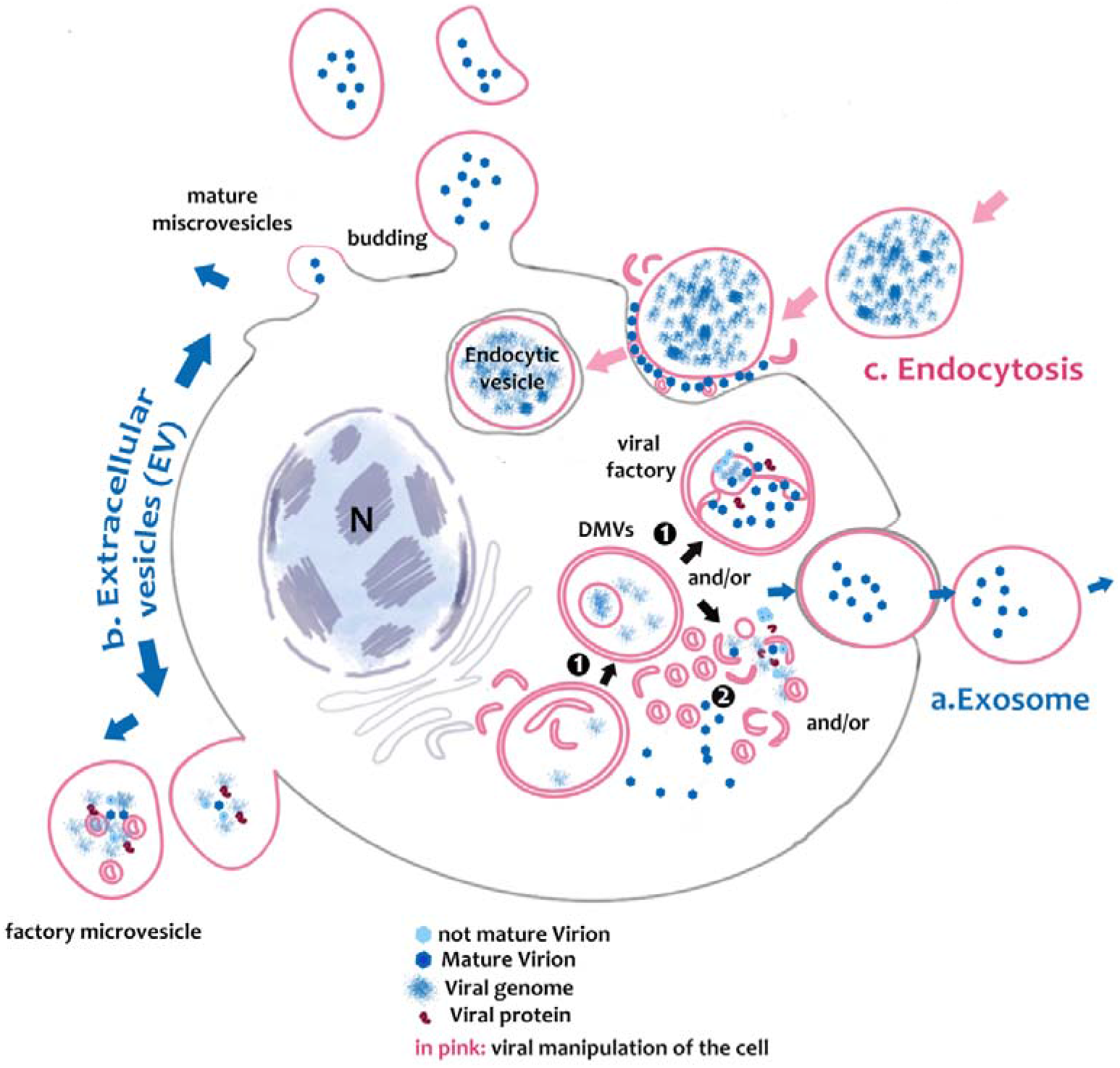
Schematic of Picornavirus spread in noble pen shell *Pinna nobilis* haemocytes. Virus can form in large DMVs (**1**), or in small microvesicles (2) where virus is freed in the cytoplasm. Then viral replication and release may be performed using different pathways. **a.** Viruses may use the autophagic-like vesicles for viral spread, packaging virions or viral factors that will be released to the extracellular medium as **exosomes** after fusion with the plasma membrane; **b.** Infectious microvesicle-like structures can either bud as evagination directly from the cytoplasm containing formed assembled virions and exit in **extracellular vesicles EVs** with mature virions or as factory-microvesicles; then **c**. PV infective mechanism involves vesicle entering through **endocytic pathway**.

**Figure 3.**
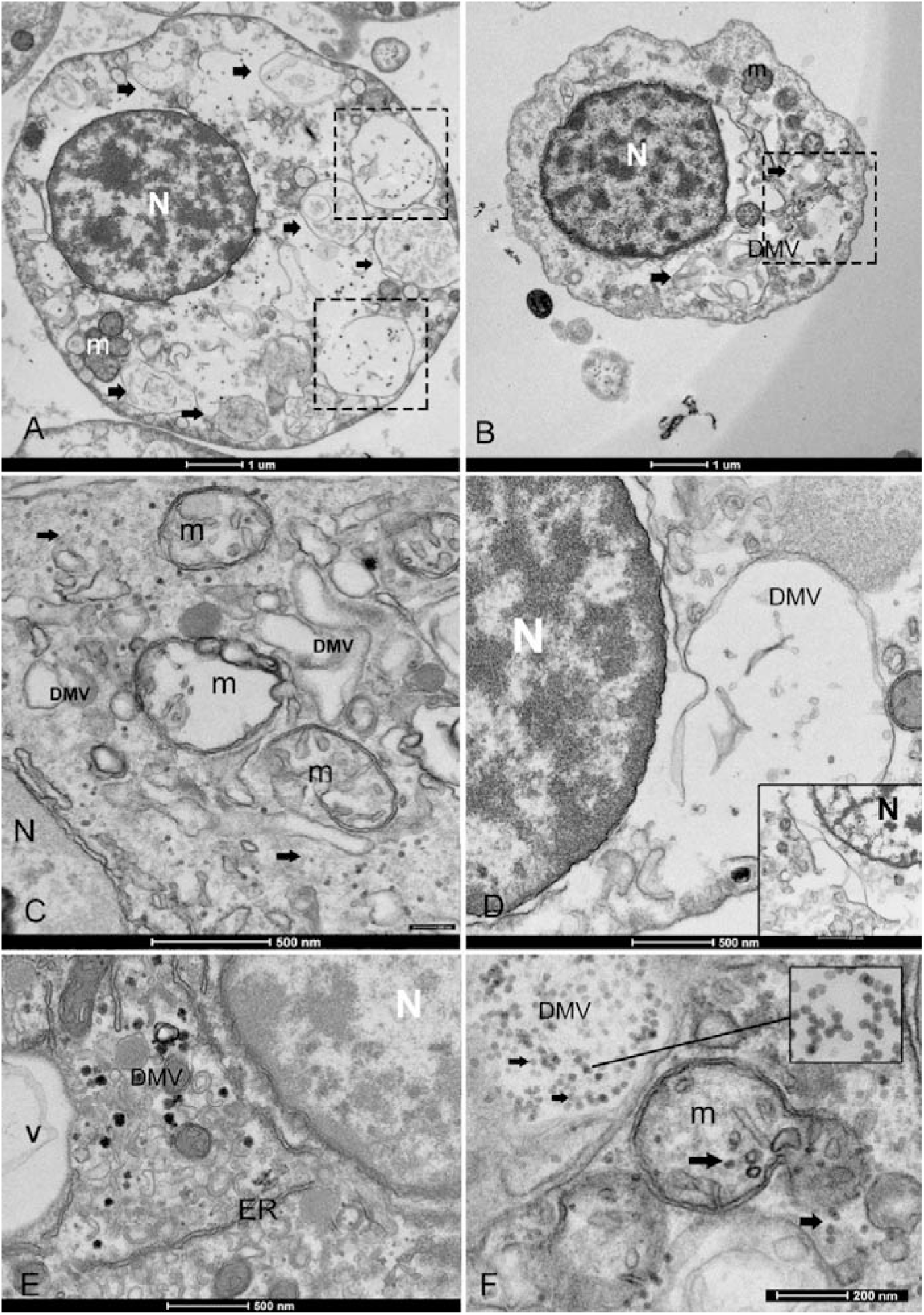
Overview of the morphological features of viral infection in the haemocytes of noble pen shell *Pinna nobilis*. **A-B.** granulocytes (**A**) and hyalinocytes (**B**) displaying double membrane vesicles (DMV) (arrows and squares). **C**. Small vesicles (100-200nm -DMVs) and damaged mitochondria (m) with visible presence of viral particles in the cytoplasm (arrows); **D.** Vesicle originates from the ER and nuclear membrane (inset); **E.** Endoplasmic Reticulum (ER) rearranged, forming DMV (v) and detail of zippered ER that consists of long stretches of ER-derived paired membranes. **F.** Detail of vesicle containing viral particles (inset), virus replication in a mitochondrion (m), but also present in the cell free in the cytoplasm (arrows). N: nucleus.

In all immune cells, aggreggates of modified membranes occupied large areas of the perinuclear cytoplasm. Numerous membranous vesicles forming virions at various stages of self-assembly were visible. Vesicles in the cytoplasm were clearly associated with a dilated ER and nuclear membrane (**Figure 3C-D**). They had different dimensions and were constituted by double membranes, reported in literature as DMVs or as *autophagosome-like* vesicles. The small vesicles measured 80-100 nm, while larger DMVs measured around 800-1000 nm (**Figure 3C-E**). Presence of viral particles related to the cytoplasmic vesicles or mithocondria membranes were visible in all immune cells of *P. nobilis* (**Figure 3F**). Smaller vesicles were mostly used for virus genome replication at cytoplasmic level, while the larger DMVs supported a later step of virus production, specifically provirion maturation and forming of complex factories. The DMVs displayed evident double or more complex membranes (**Figure 4A**). In earlier phases, they were filled by an unassembled viral amorphous and granular material; vesicles then showed one or two ring-*like* structures captured in the lumen vesicle, as also reported in other PVs (Yang et al., 2020) (**Figure 4B**). Viral replication factories were typically constituted by complex membrane remodelling emerging from the DMVs formed by viral particles and proteins assembling mature virions, in some cases packed together. The virion is a an icosahedral, non enveloped, small (20 nm) particle with no discernible projections (**Figure 4C-D**).

**Figure 4.**
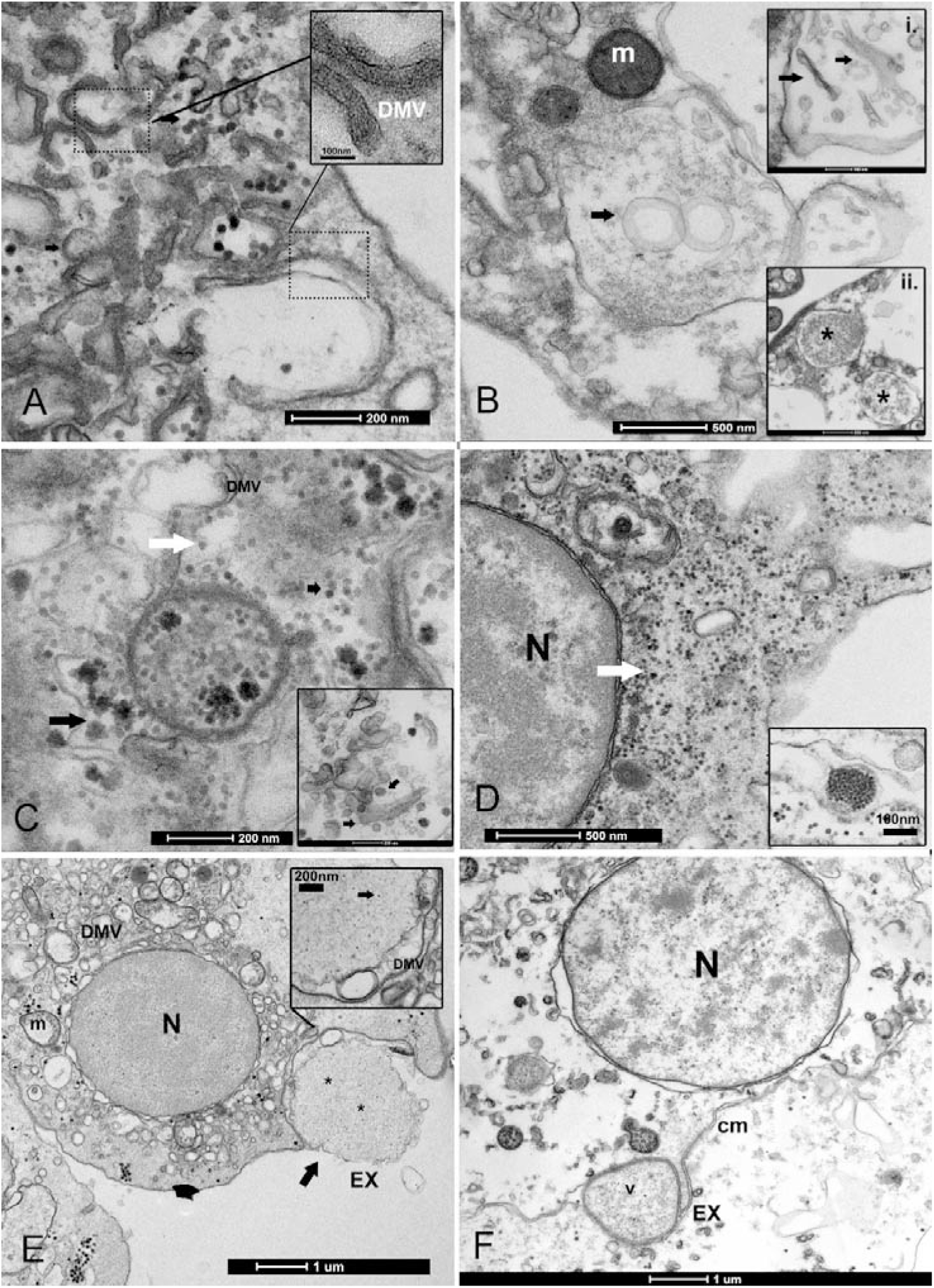
Biogenesis of Double Membrane Vesicles (DMV) and exosome vesicles during PV replication. **A.** details of small and big DMVs densely packed mebranes that have an active role in viral RNA synthesis. **B.** DMV biogenesis requires several membrane-remodeling steps: the biogenesis of DMVs can occur by the induction of positive membrane curvature, through membrane pairing (inset i); these structures form cisternae that curve to finally seal and transform into a closed DMV and forming ring like vesicles as reported in other picornavirus where viral material can assemble (inset ii); **C-D.** the virus replicates into factories through remodeled double membranes (inset) producing mature virions (white arrow) accumulating in these large intracellular compartments from immature and forming ones (black arrow). **D.** PV spreading into the granulocyte cytoplasm (white arrow); inset: grouped mature virions into a vesicles. **E-F**. Vesicle filled with viral particles (*) released to the extracellular medium as exosomes (EX) after fusion with the cell membrane; the viral egress involves the fusion of a double-membrane vesicle (V) with the cell membrane (CM); N: nucleus; m: mitochondria.

Once the virus was assembled, viral release could occur through different mechanisms.Viruses could be released without lysis, through the so-called secretory autophagy, after the fusion of the DMVs with the plasma membrane, and released-as exosomes. Exosomes typical dimension ranged between 800-1000 nm. They were secreted after their fusion with cell membrane vesicles and released as single-membraned virus-filled vesicles (**Figure 4E-F**). The 4 samples from Venice lagoon, defined in the field as apparently healthy, displayed mature viral particles spreading into cytoplasm (**Figure 4D**), small DMVs and active factories.

The predominant infective strategy observed in PV of *P. nobilis*, typical for non-enveloped viruses, was a non-lytical “unconventional secretion” using extracellular vesicles (EVs) (**Figure 5**). PV-infected cells secreted EVs carrying viral RNA particles. Vesicles ranged in a diameter from 100 to 700 nm. Virions may be released from cells by a budding process from both granulocytes and hyalinocytes (**Figure 5A-C**). Infective vesicles were seen carrying either mature viral particles (**Figure 5D**) or vesicle factories - containing still immature virions and DMVs (**Figure 5E-F**). EVs may be taken up by recipient cells by endocytosis or fusing with the plasma membrane. Through endocytosis, the vesicle is internalized in an endocytic vacuole surrounded by DMVs to form then mature viral particles (**Figure 6A-B**). Infective vesicles transporting mature virions injecting their contents were also visible (**Figure 6 C-D**).

**Figure 5.**
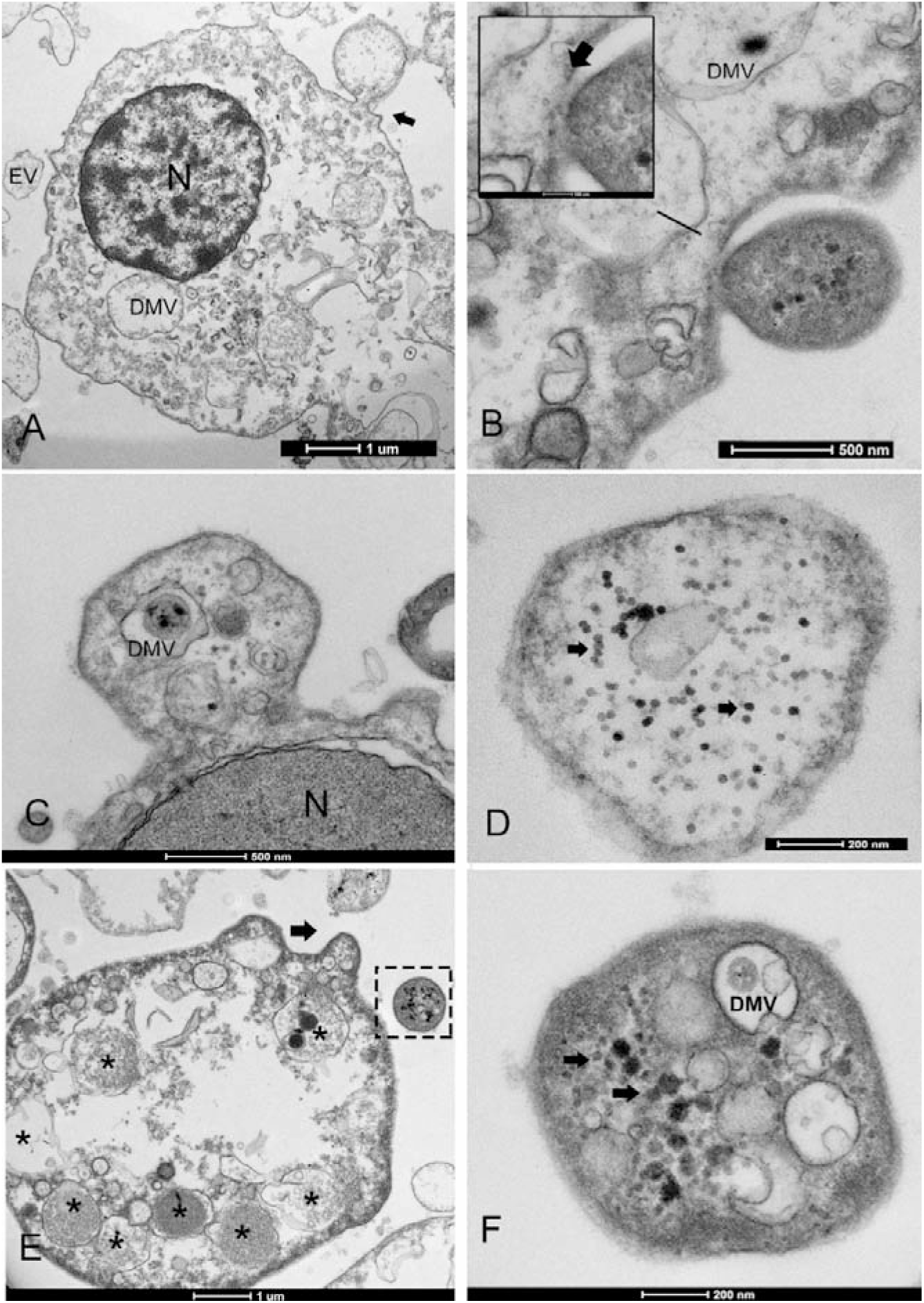
Extracellular vesicle (EV) formation during PV infection. **A-B.** The EV lipid membrane mediate the release of virions through budding formation (arrow) with shedding microvesicles, **B inset**: detail of the budding from the cell membrane (arrow). **C.** Hyaloncyte budding formation; **D.** Vesicles might contain mature virions (arrows) or viral and/or host factors viral components and induced/altered host factors (**E-F**); N: nucleus; DMV: double membrane vesicles within the EV.

**Figure 6.**
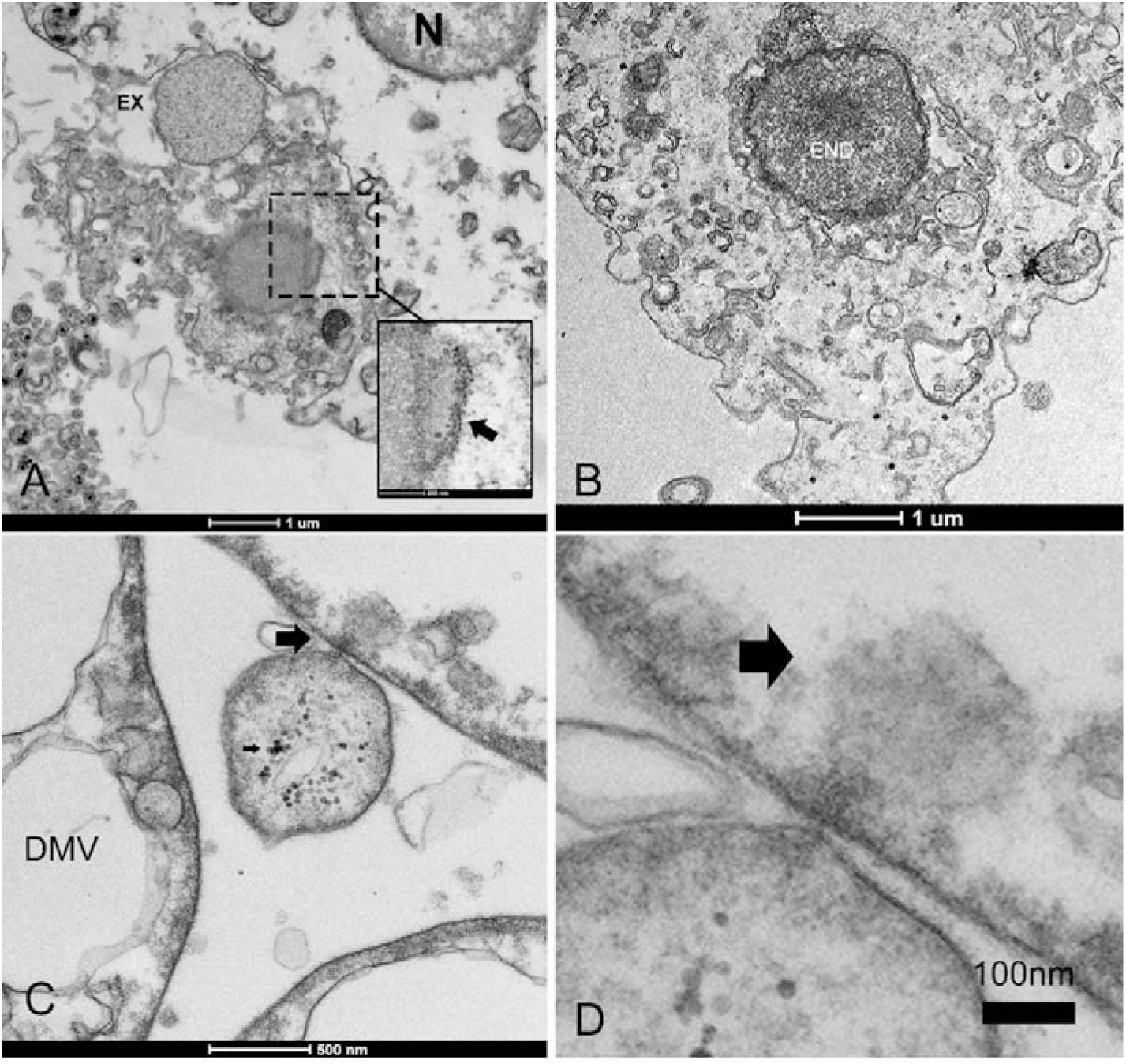
Infectious vesicles containing viral particles entering the cells. **A-B.** Infective vesicle containing unassembled viral particles can enter through endocytosis fusing with cellular membrane and forming virions (inset, arrow) during a release of Exosome (EX), and then transported as late endosomes (END) close to the nucleus (**B**); **C-D**. Vesicle can inject its content into other cells (arrows).

Dead haemocytes were observed during infections. The highest number of damaged cells was observed in animals maintained in captivity at IMEDMAR-UCV and Murcia Aquarium, surrounded by an elevated number of EVs or containing endocytic vesicles. Damaged cells appeared empty, they were shrumk and contained few vesicles, and presented membrane rupture and apoptotic bodies (**Figure 7**).

**Figure 7.**
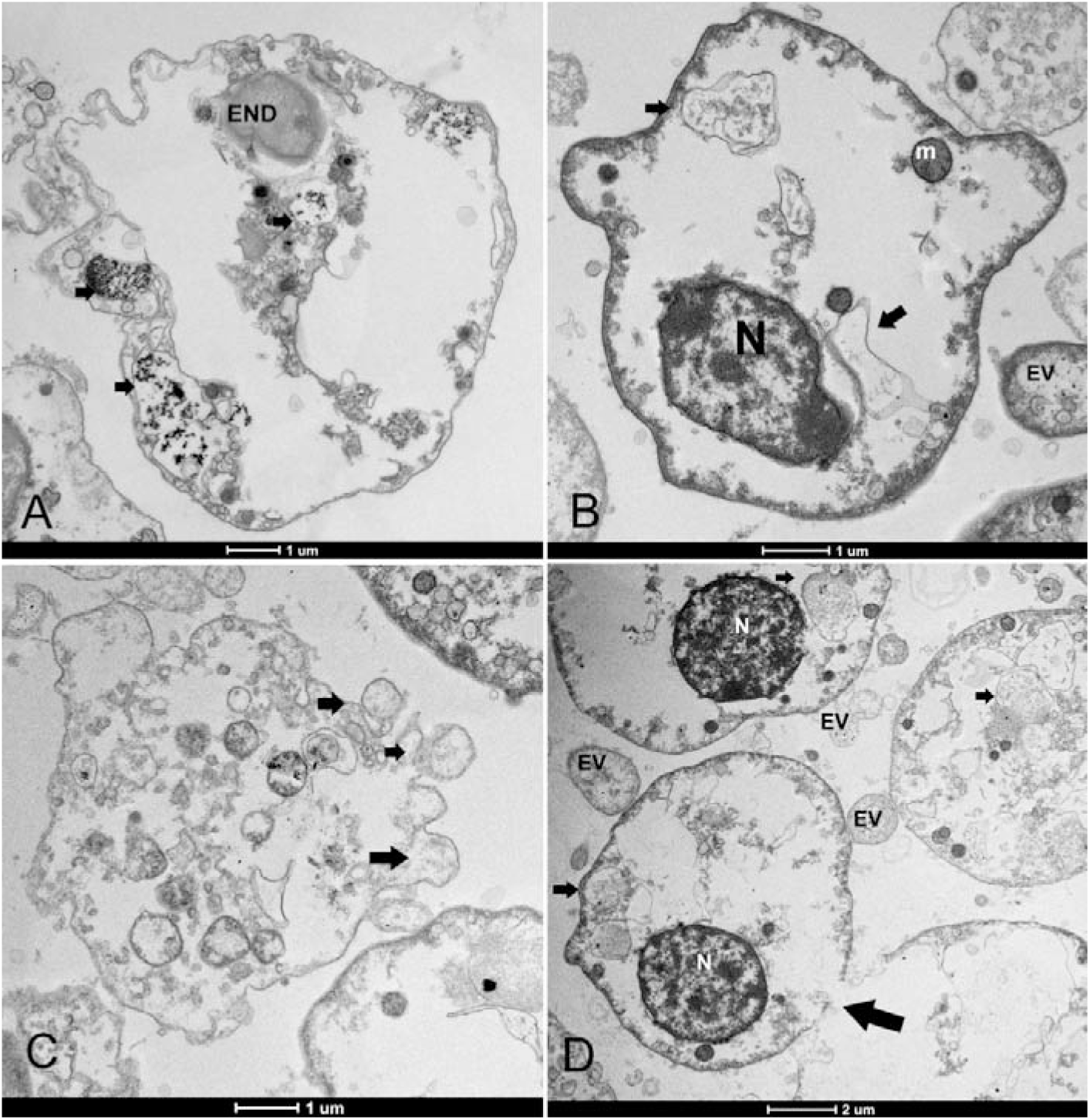
Haemocytes cell death. **A-B.** Changes mainly include empty cytoplasm with residual vesicles (arrows) or endocytotic vesicle (END) sourrounded by extracellular vesciles (EV); **C.** blebbing of the plasma membrane and formation of apoptotic bodies (arrows) and loss of nuclei; **D.** membrane rupture (**arrows**). N: nucleus; m: mitochondria.

### 3.2 Identification of RNA viruses in *P. nobilis* haemocytes

Sequencing of six stranded cDNA libraries retrieved from haemocyte samples of six *P. nobilis* using Illumina technology generated a total of 162,139,300 paired-end raw reads (**Table 2**), subsequently deposited in the NCBI Sequence Read Archive (SRA) under the BioProject accession PRJNA999583. After trimming and mapping against the *P. nobilis* genome, the number of unmapped paired reads was 33,097,808 (20.4%). These unmapped reads were used to assemble the viral RNA genomes present in the *P. nobilis* haemocytes.

BLASTX analysis of the 140,922 contigs assembled using Trinity resulted in 61,634 positive matches (43.7%, e-value < 10-20) against the viral protein database, encompassing DNA and RNA viruses, as well as bacteriophages. Since this study aimed to identify RNA viruses in the *P. nobilis* haemocytes, we focused specifically on this viral group, usingVirsorter2 and VirBot software tools. Both analyses identified the same set of five contigs, ranging from 5210 to 8876 nucleotides (nt) in length, all classified, with a score of 1 (the highest possible score with Virsorter2), into the realm *Riboviria*.

These five sequences exhibited the highest BLASTX matches with different *Riboviria* polyproteins 1 and 2 (**Table 5**), mainly of *Picornavirales*. Notably, two of these contigs (TRINITY_DN35348_c1_g1 and TRINITY_DN9545_c0_g1) displayed the highest number of mapped reads, particularly in the Trab samples (Site 1, Trabucador), indicating their abundance among the five identified viruses. Additionally, during the global BLASTX analysis, we discovered the contig TRINITY_DN58105_c0_g1_i1 (841 nt), that matched with the predicted replication-associated protein of *Chaetoceros tenuissimus* RNA virus type II (BAP99820.1). Manual overlapping with TRINITY_DN35348_c1_g1 (5210 nt, matched with the predicted structural protein of *Chaetoceros tenuissimus* RNA virus type II, BAP99821.1), resulting in a 6023 nt consensus sequence.

**Table 5.**
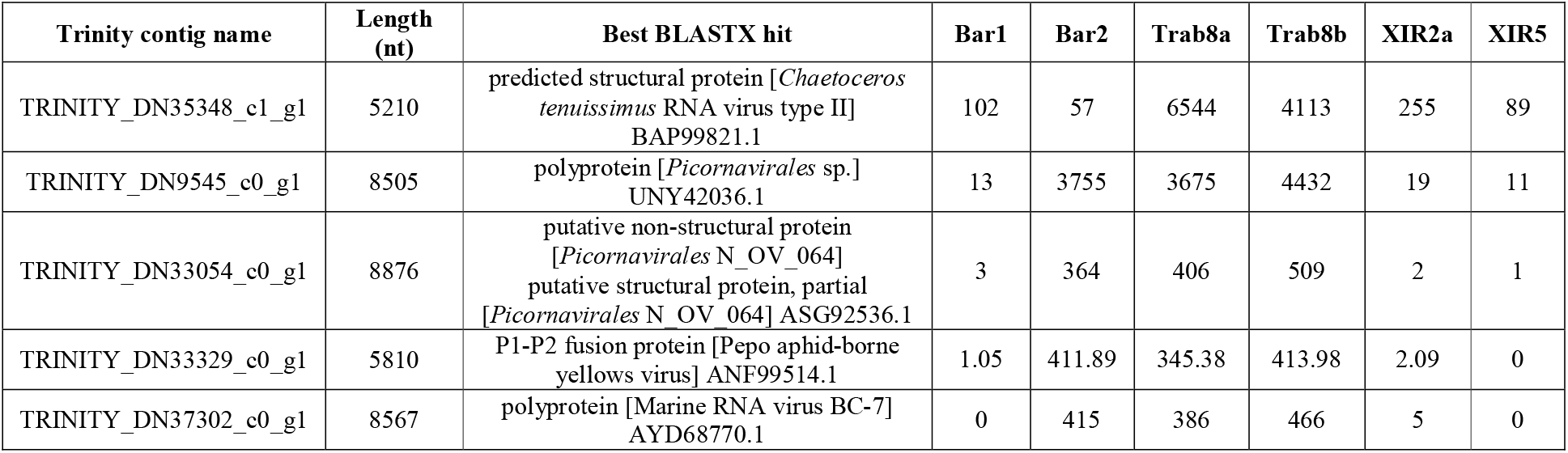
Assembled contigs classified into the realm *Riboviria* by Virsorter2 and VirBot analysis, results of their annotation through BLASTX versus the viral protein database and reads abundance in the different noble pen shell *Pinna nobilis* samples expressed as raw counts.

**Figure 8** presents the genomic organization of the two most prevalent viruses identified herein. The sequence consensus_DN35348_c1_g1_DN58105_c0_g1 (A) is partial, lacking the complete 5’-end; while the DN9545_c0_g1 sequence (B) is complete. These two sequences were deposited in GenBank under accessions OR448788 and OR448789. The ORFfinder analysis identified two large ORFs, encoding two viral polyproteins, in both sequences. Within the polyproteins 1, there were the conserved domains of RdRp and RNA helicase (this latter only detected in sequence DN9545). In contrast, structural polyproteins 2 have two rhv-like domains (picornavirus capsid protein domain-like), a CRPV capsid domain (a family of capsid proteins found in positive stranded ssRNA viruses), and a short VP4 domain (a family of minor capsid proteins, located within the viral capsid, at the interface between the external protein shell and packaged RNA genome). The presence of two large ORFs encoding these specific domains suggests that these viral sequences belong to the *Picornavirales* order.

**Figure 8.**
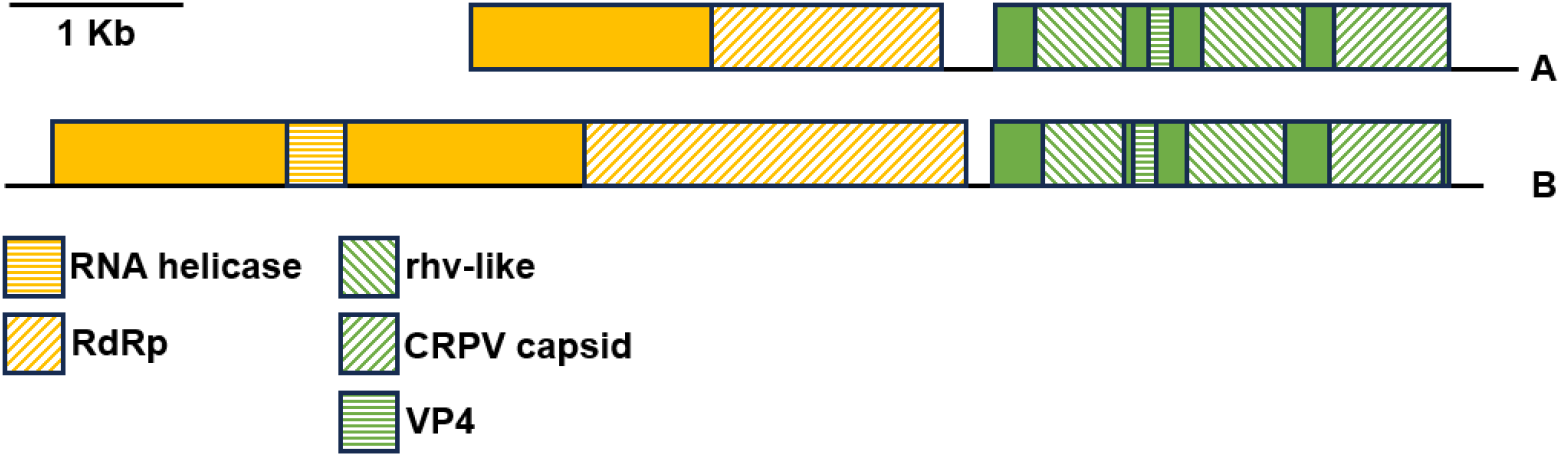
Genomic organization of the most abundant RNA viruses identified in the noble pen shell *Pinna nobilis* haemocytes. The colored boxes represent the ORFs of the genes 1 (yellow) and 2 (green) encoding for the viral polyprotein 1 and 2, respectively. The different colored motifs represent the regions encoding specific viral proteins. A) Trinity assembled contigs DN35348_c1_g1 and DN58105_c0_g1, 5’-end partial (accession number OR448788); B) Trinity assembled contig DN9545_c0_g1 (accession number OR448789).

The genomic organization of the remaining three viruses identified in *P.nobilis* haemocytes found herein (DN33054, DN37302, and DN33329) were deposited in GenBank under accession numbers OR448790, OR448791, and OR448792, respectively (**Supplementary** Figure 1). Similarly to the two most abundant viruses, the genomic organization of DN33054 and DN37302 presented two large ORFs, with ORF1 encoding RdRp and RNA helicase, and ORF2 encoding two rhv-like domains -a CRPV capsid domain, and a short VP4 domain. In contrast, the genomic organization of DN33329 was quite different, with three partially overlapping main ORFs encoding a peptidase and the RdRp, similar to the structure of the *Sobelivirales* order.

The Maximum Likelihood tree was generated based on the alignement of the conserved region of the RdRp proteins of the viruses identified in the *P. nobilis* haemocytes with homologous sequences downloaded from GenBank (**Figure 9**). Almost all of the RdRp sequences downloaded from GenBank belonged to *Picornavirales* from marine or freshwater environmental samples. A restricted number of sequences belonged to *Sobelivirales*, and two of them (*Penaeus vannamei* picornavirus and Wenzhou shrimp virus 8) to *Dicistoviridae* that are associated with mollusc and crustacean diseases (Liu et al. 2021; Srisala et al., 2023). Amongst the viral sequences identified in *P. nobilis* haemocytes, DN33329 clustered with those in the order *Sobelivirales*, along with viruses from rivers, soil, and plants, agreeing with the genomic organization previously described (**Supplementary** Figure 1). Also The remaining four sequences from *P. nobilis* belonged to the order *Picornavirales*, as expected from their genomic organization; among them, the two most abundant are clustered within the family *Marnaviridae*.

**Figure 9.**
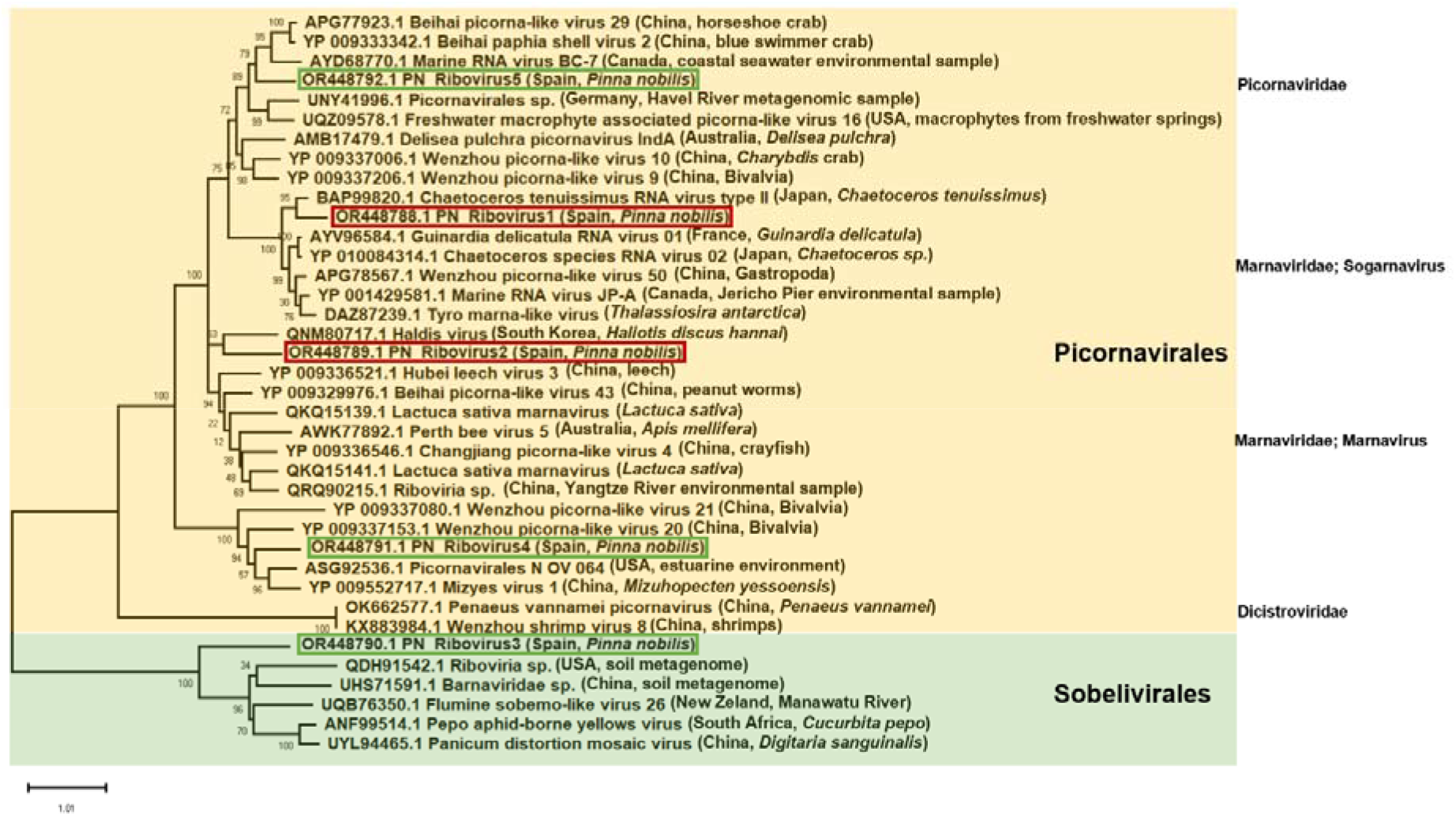
Maximum Likelihood tree constructed using the amino acid alignment of the conserved region of the RpRd of the RNA viruses identified in the noble pen shell *Pinna nobilis* haemocytes and other RpRd proteins downloaded from Genbank (total 38 amino acid sequences). The tree with the highest log likelihood (-16083.36) is shown. The tree is drawn to scale, with branch lengths measured in the number of substitutions per site. The red and green boxes highlight the most and less abundant RNA viruses identified, respectively. The bootstrap percentages inferred from 500 replicates are shown next to the branches. When available, the geographic origin and host of the samples are indicated within parentheses.

In term of ultrastructure, both samples from the Trabucador area (Trab8a and Trab8b) that had highest number of mapped viral reads displayed numerous cytoplasmic viral factories and extracellular infective vesicles spreading into the hemolymph, suggesting an active infective phase.

### 3.3 Detection of RNA virus in *P. nobilis* haemocytes by quantitative real-time PCR

The quantitative real-time PCR experiments conducted on the cDNA of the *P. nobilis* haemocytes collected in different sampling points in Catalonia were positive for the primer pair PNChetoF/R designed on the reconstructed DN35348_DN58105 sequence, similar to a *Chaetoceros tenuissimus* RNA virus type II. These results confirm the *in silico* analysis of viral reads abundance (**Table 5**), with Trab8a and XIR2a showing the highest infection level and XIR5 showing a low infection level (**Figure 10**). With the same primer pair, viral RNA was detected in all other examined samples presenting variable infection levels. The primer pair PNPicorF/R designed on the sequence DN9545 did not provide detectable amplification signal.

**Figure 10.**
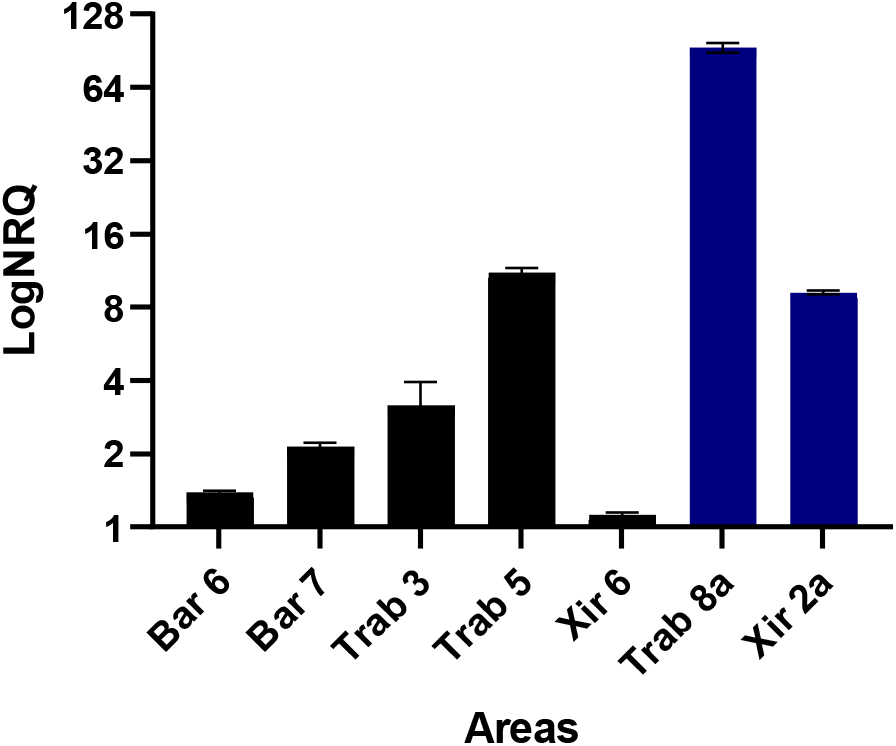
Results of the qPCR experiment conducted on the cDNA of the noble pen shell *Pinna nobilis* haemocytes collected in different areas using primer pairs specific for the *in silico* assembled Sogarnavirus (accession number OR448788). Bars represent the logarithm of the mean Normalized Relative Quantification ± SEM using the *P. nobilis* actin as endogenous gene. The blue bars represent the samples used for the *in silico* RNA-seq experiment.

## 4. Discussion

This study reports the discovery of a picornavirus affecting *P. nobilis* haemocytes in Spain and Italy during MMEs (Carella et al., 2020; Saric et al. 2020; Carella et al., 2019). Previous studies investigating the etiological agent involved in the MMEs showed a complex pattern: disease diagnostic repeatedly confirmed the simultaneous presence of several agents like *Haplosporidium pinnae*, *Mycobacterium* spp. (Carella et al 2019, 2020, 2023a), *Vibrio mediterranei* (Prado et al., 2020), and in some cases *Perkinsus* spp., suggesting that a common primary cause, not yet identified, could be linked to the phenomena (Carella et al., 2020). Based on our observations, this picornavirus could be the underlying cause of these events, once the virus was actively replicating and infecting the *P. nobilis* shell immune cells in 100% of the examined residual populations in Italy and Spain, from 2021 to 2023. In fact, the detection of this virus in the same populations over a 2-year period suggests that animals can maintain persistent infections until the disease becomes chronic, weakening them, and becoming susceptible to opportunistic infections, as also reported in other immunosuppressive viruses (English et al., 1994). This could explain the apparent resistance of the residual individuals that are slowly dying in all areas. The affected animals remain apparently healthy for months to years, before the immune system begins to collapse, as reported by many scientists (Carella et al., 2023a; Saric et al., 2020; Zotou et al., 2020).

In this study, the observed picornavirus affecting *P. nobilis* in Italy and Spain used granulocytes and hyalinocytes as vessels for dissemination, replication and long-term persistence, interfering with haemocyte immune abilities, as observed in other picornaviruses (Cusik et al., 2014). Multiple known immunoregulatory mechanisms play a role in persistent viral infections, resulting in immunosuppression (Oldstone, 2009). Immunodeficiency viruses have been reported worldwide in felid, bovine (dairy cattle), and primates (chimpanzees) species, and have been observed actively replicating in the blood and lymphoid tissues (Tizard, 2019). As a result of complete or unsuccessful viral replication, immunodeficiency viruses may functionally lyse or impair immune cells like lymphocytes, as observed in parvovirus infections of B-lymphocytes, in infectious mononucleosis due to Epstein-Barr virus, and of CD4+ T-lymphocytes and macrophages in AIDs caused by HIV lentivirus. (Ackley et al., 1990; Brown et al., 2010). In other cases, viruses can infect and damage cells involved in phagocytosis, like macrophages, as seen in influenza virus and dengue virus infections (Egberink and Horzinek, 1992; VandeWoude and Apetrei, 2006). Similarly, PV can block the production of type I interferon (IFN), leading to T cell depletion, or can modulate immune cells apoptosis and autophagy mechanisms (Lei et al., 2011; Lee et al., 2012; Son et al., 2013). In bivalves, the primary role of haemocytes is pathogen killing and elimination through phagocytosis, encapsulation, production of cytotoxic molecules, antimicrobial peptides, and secretion of inflammatory cytokines (de la Ballina, 2022; Song et al., 2010; Cheng, 1981). A morpho-functional description of *P. nobilis* haemocytes was performed prior to the mortality events in 2015, showing that healthy haemocytes were able to conduct phagocytosis (Matozzo et al., 2016). In our study, *P. nobilis* haemocytes from individuals sampled in Spain and Italy displayed damage in mitochondria, ER and other organelles, which may have affected their regular immune function (Pugliese et al., 2005). This is also true considering that a preliminary study perfomed in *P. nobilis* shell haemocytes from the same areas, in 2021-2022, already revealed strong immunodepression, represented by decreased immune cells response to pathogenic stimuli following an *in vivo* challenge (Carella et al., 2023b).

Immunosuppression is necessary for the PVs to use cell components in order to promote infection (Cusik et al., 2014). PVs have evolved the ability to usurp the infected cells’ endogenous functions, inducing (endo)membrane rearrangements that are referred to as viral replication organelles (Meckes et al., 2011). Recent work revealed that picornaviruses pack multiple viral particles into a cellular microvesicle, to promote the spread of several heterogeneous virions at a time (Chen et al. 2015). In this context, the production of EVs and endocytic vesicles play an important role in the pathogenesis and in establishment of viral genome persistence and latency; moreover, viruses that establish chronic infections, have been shown to modulate the production and content of more infectious EVs, increasing their infective potential (Amari and Germain, 2021). Animals maintained in captivity in this study, displayed a high number of infective vesicles and dead cells, also related to hemocytopenia. PV persistent infection can induce lymphopenia in human and animals, like cows, cats and birds, as a direct result of viral infection, or indirectly, through cytokine induction (Cusik et al., 2014). Along with other diagnostic indices, decreased THC is a critical condition in the progression of an infection or a disease in invertebrates (Cheng et al., 2004). Heamocytopenia has been observed during chronic viral infections like in abalone ganglioneuritis (Hooper et al., 2007), and during bacterial and parasitic diseases in other mollusc species (Shields et al., 1997; Handlinger et al., 2004). Typically, persistent infections bring to a relatively small pool of haemocytes into the haemolymph sinuses and perivisceral beds, and such sequestration into tissues may lead to a dramatic decrease in haemocyte numbers (Hooper et al., 2012. before the observation mass mortality in the Mediterranean, Matozzo, (2016), reported a higher number of circulating haemocytes (5x10^5^ cells) in comparison to the current situation described in Spanish wild and captive populations. Based on our results, this could indicate an over time immune cell decrease due to infection, and a worsen situation for animals maintained in tanks. Apart from haemocyte sequestration into infected tissues, such heamocytopenia is also possibly caused by apoptosis due to haemocyte infection (Sun et al., 2019). This is a host-initiated process of programmed cell death essential for normal cell turnover, that also functions as an antiviral mechanism to limit the spread of a virus (Below et al., 2003; Croft et al., 2017).

In the present study, we were able to assemble different *Riboviria* sequences of the orders *Picornavirales* and *Sobelivirales*, with NGS Illumina sequencing obtaining nearly complete five different genome accompanied by a phylogenetic analysis. Our findings support previous observations that bivalve and other invertebrate viromes harbor a broad diversity of viruses of the order *Picornavirales* (Shi et al., 2016; Zhou et al., 2022), matching with the ultrastructural feature of haemocyte PV infecting. Picornavirus are components of the virome associated with filter-feeding organisms under normal physiological conditions, reported in metatranscriptomic studies in absence of detectable disease (Moreira et al., 2012). Among the five assembled *Riboviria*, the two most abundant sequences clustered with the *Picornavirales* group within the family *Marnaviridae*, which comprises small non-enveloped viruses that infect various photosynthetic marine protists (Sadeghi et al., 2021), and also reported in diatom population (Shirai et al., 2008). The most abundant reads placed the haemocyte PV close to a virus isolated in the marine diatoms *Chaetoceros tenuissimus*, named *C. tenuissimus* RNA virus type II (Shirai et al., 2008), belonging to the genus *Sogarnavirus*; while the second most abundant PV was identified as *Picornavirus* belong to the genus *Marnavirus*.

Microalgal cells infected by *Marnaviridae*, genus *Sogarnavirus*, can display extensive cytopathic effects with ultrastructural changes, including swelling of the endoplasmic reticulum, vacuolation and disintegration of the cytoplasm (Nagasaki et al., 2004; Shirai et al., 2008), as reported in our study. To our best knowledge, this is the first report of a sogarnavirus affecting an animal species. In order to confirm the NGS analysis, a qPCR was perfomed to amplify the most represented *Sogarnavirus*. The primer pairs designed on the sequence of the *P. nobilis* picornavirus similar to the *C. tenuissimus* RNA virus type II were positive in the analysed samples; while no amplification was obtained with the other primer pairs. These results may not be considered conclusive and need to be confirmed with other analysis.

In this study, both apparently healthy and sick animals showed the ultrastructural features of viral infection. In confined waters across Mediterranean regions, like lagoons, MMEs started later (Saric et al., 2020) For intance, in Venice lagoon the mortality episodes began in 2020, later to the *P. nobilis* populations die offs of the Ebro Delta, which started in 2018 (Carella et al., 2020). The viral infection could have originated first in Spanish waters, years before mortalities became evident, subsequently reaching other areas, until animals presented an acquired immunodeficiency (i.e., low count of haemocytes), as occurs with other immunodeficency viruses. The disease, made of periods of remission followed by disease progression over time, ends up gradually weakening the populations that start display symptoms, leading to decline. These individuals are exposed to a wide variety of bacterial and parasitic opportunistic infections, acting in an unfavorable situation. In the Venice lagoon, in the area where animals appeared apparently healthy in May 2023, a remarkable mortality was observed two months later. In the 4 analyzed individuals, all immune cells presented actively replicating virus. This means that infection can be subclinical until the very late, more likely when an opportunistic pathogen takes advantage of the situation.

As previously reported, recent findings suggested a more complex scenario than the one originally proposed in the first descriptions of *P. nobilis* MMEs (Carella et al., 2019, 2020, 2023a). Diseases with complex etiology are influenced by various interacting drivers, and frequently remain difficult to characterise. Indeed, this viral disease represents an epidemiologic threat of unknown proportions for marine animal population. Further studies are required to determine the ecological importance of PV as an agent affecting the dynamics of *P. nobilis* MMEs in the analyzed areas and in other natural environments.

## 5. Conclusions

Few viruses have been associated with disease outbreaks and mortality in bivalves, belonging mainly to the families of *Alloherpesviridae* and *Iridoviridae*. Nevertheless, picornavirus-*like* particles have also been reported during disease outcome (Carballal et al., 2003; Rasmussen, 1996; Ip and Desser, 1984). Here we report the first description of an immunodeficiency virus in bivalves, affecting *P. nobilis* haemocytes. Immune defence is a key determinant of fitness underlying the capacity of animals to resist or tolerate potential infections. Immune function is not static and can be suppressed, depressed, or stimulated by exposure to environmental abiotic factors, food availability and pathogens. Alteration of this response can directly affect population dynamics and its survival. Lately, the emergence and spread of new and existing pathogens pose a significant risk not only to human life and animal health, but also to the conservation of wildlife. Future studies are needed to establish a clear and significant correlation between the genome of the picornaviruses assembled in *P. nobilis* haemocytes and the *P. nobilis* MMEs. Firstly, the full length of the partial PV sequence assembled *in vitro* should be obtained using a 5’RACE approach. Secondly, viral isolation and cultivation attempts are required to define virus charactheristics. Finally, a fast and reliable diagnostic test (e.g., based on qPCR) should be develop in order to rapidly check the infection level of the surviving *P. nobilis* populations in other geaographic areas. Virus presence should also be evaluated in the environment (water, plankton, sediments) and surrounding animals, which could be potential virus reservoirs.

## Acknowledgements

The research is partially supported by the EU LIFE Programme Project “Protection and restoration of *Pinna nobilis* populations as a response to the catastrophic pandemic started in 2016” (LIFE PINNARCA) [grant number LIFE20 NAT/ES/001265]; the authors thank the Dr. Sergio Sorbo of the CeSMA, section of Microscopy LaMMEC, University of Naples Federico II and Dr. Antonella Giarra, Chemistry Department of University of Naples Federico II for the technical support in the electron microscopy. Authors would like to thank José Luis Costa and David Mateu Vilella (IRTA), Roberto Vianello with Andrea Sabino (CNR-ISMAR) for technical assistance during fieldwork sampling in Catalonia and Venice, respectively.

---

**Supplementary Figure 1.**
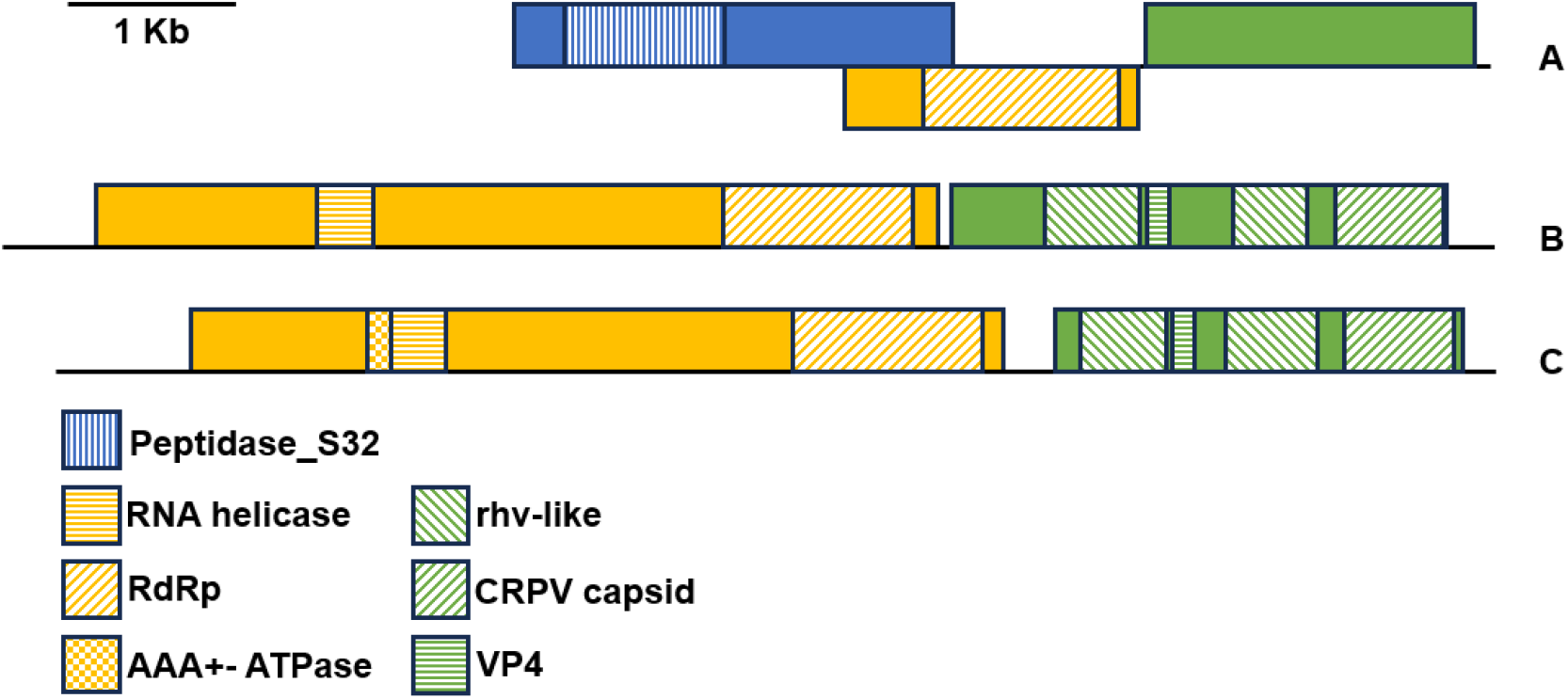
Genomic organization of the less abundant RNA viruses identified in the noble pen shell *Pinna nobilis* haemocytes. The colored boxes represent the ORFs of the genes 1 (yellow) and 2 (green) encoding for the viral polyprotein 1 and 2, respectively A) Trinity assembled contig DN33329_c0_g1 (accession number OR448790); B) Trinity assembled contig DN33054_c0_g1 (accession number OR448791); C) Trinity assembled contig DN37302_c0_g1 (accession number OR448792).

