## Supplementary figures and images for "A WIDESPREAD PICORNAVIRUS AFFECTS THE HAEMOCYTES OF THE NOBLE PEN SHELL (*PINNA NOBILIS*) LEADING TO IMMUNOSUPPRESSION"

### Supplementary_Figure 1

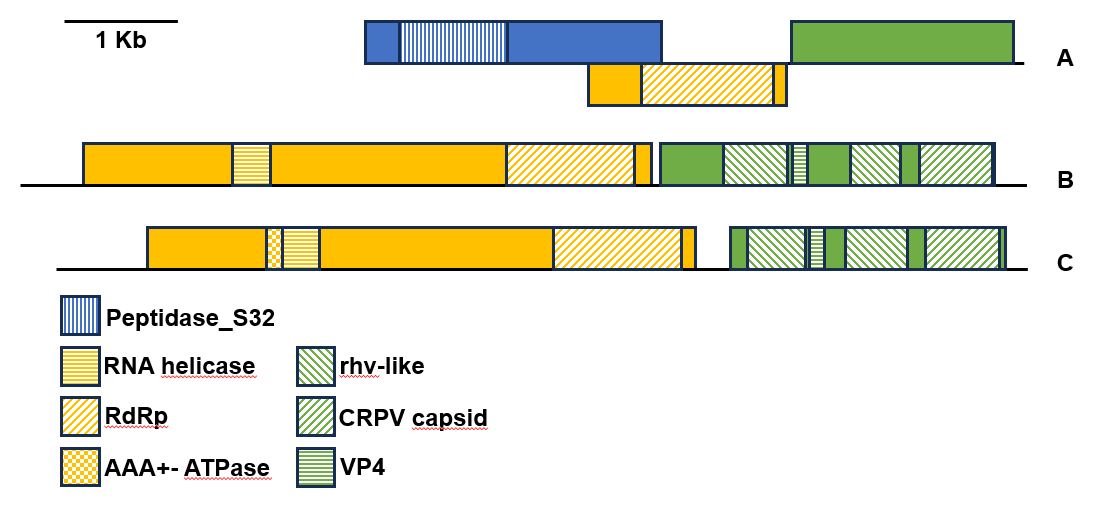
